## Supplementary figures and images for "Empirical single-cell tracking and cell-fate simulation reveal dual roles of p53 in tumor suppression"

### Figure 1

Figure 1

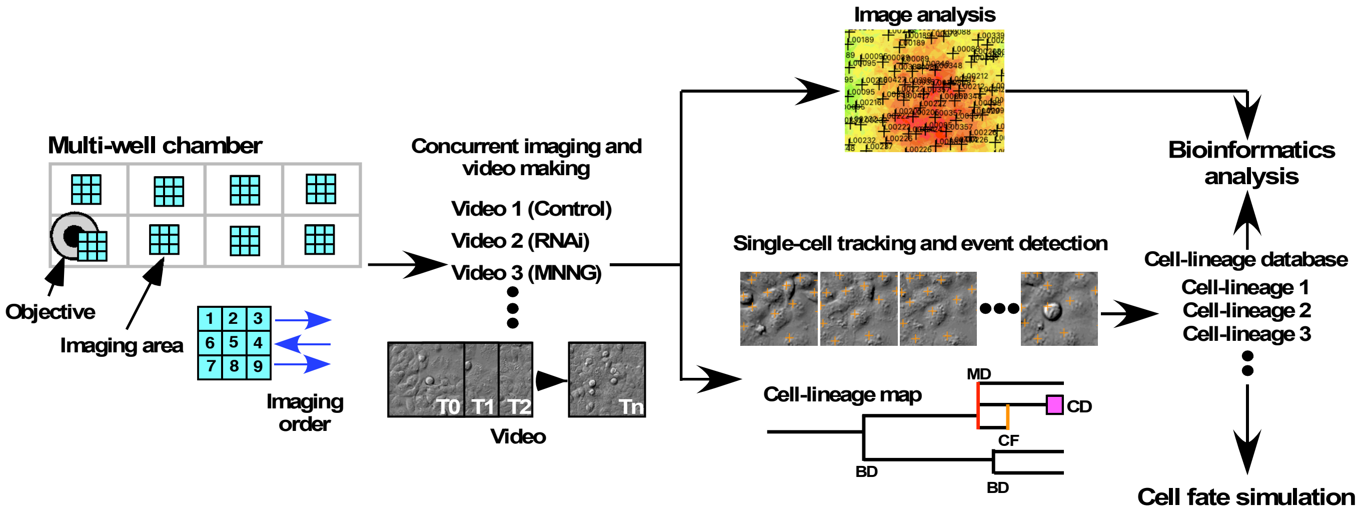


Rancourt et al.

### Figure 1-figure suplement 1

#### Figure 1-figure supplement 1

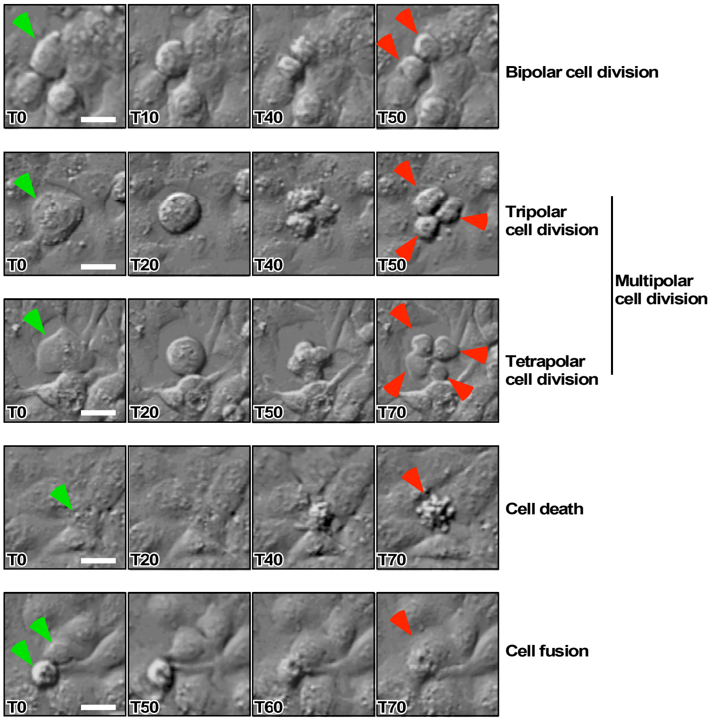


Rancourt et al.

### Figure 1-figure suplement 2

#### Figure 1-figure supplement 2

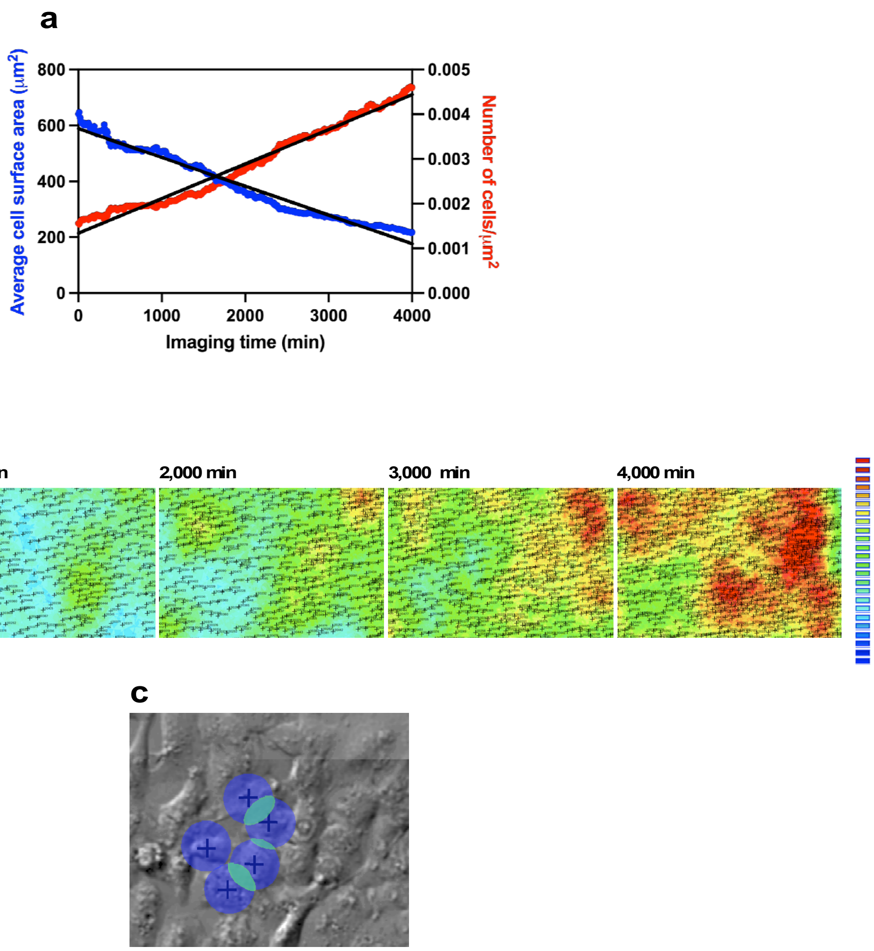


Rancourt et al.

### Figure 1-figure suplement 3

#### Figure 1-figure supplement 3

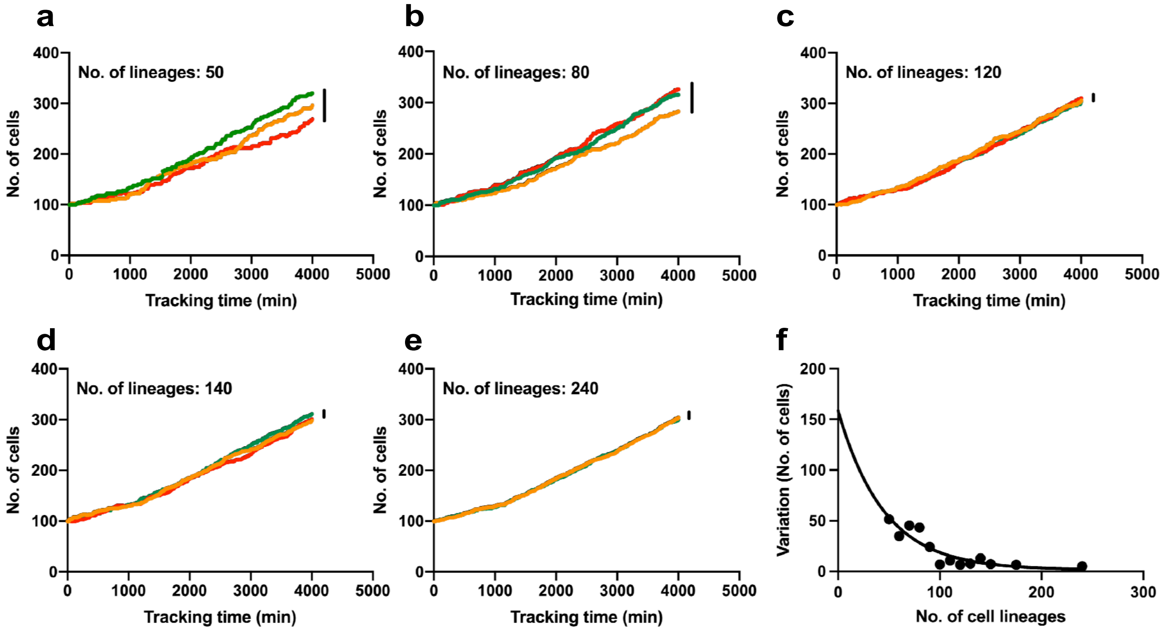


Rancourt et al.

### Figure 1-figure suplement 4

##### Figure 1-figure supplement 4

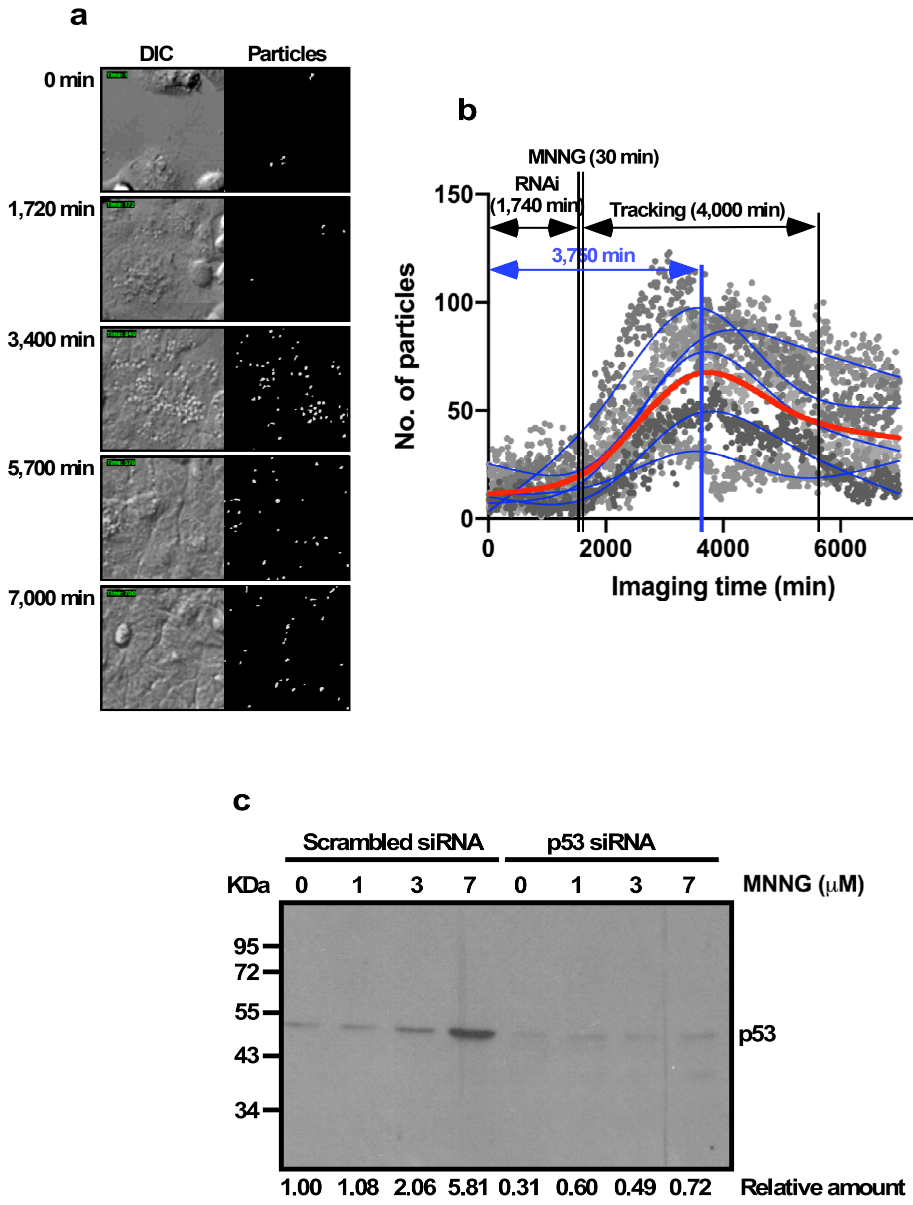


Rancourt et al.

### Figure 2

#### Figure 2

Cell counting

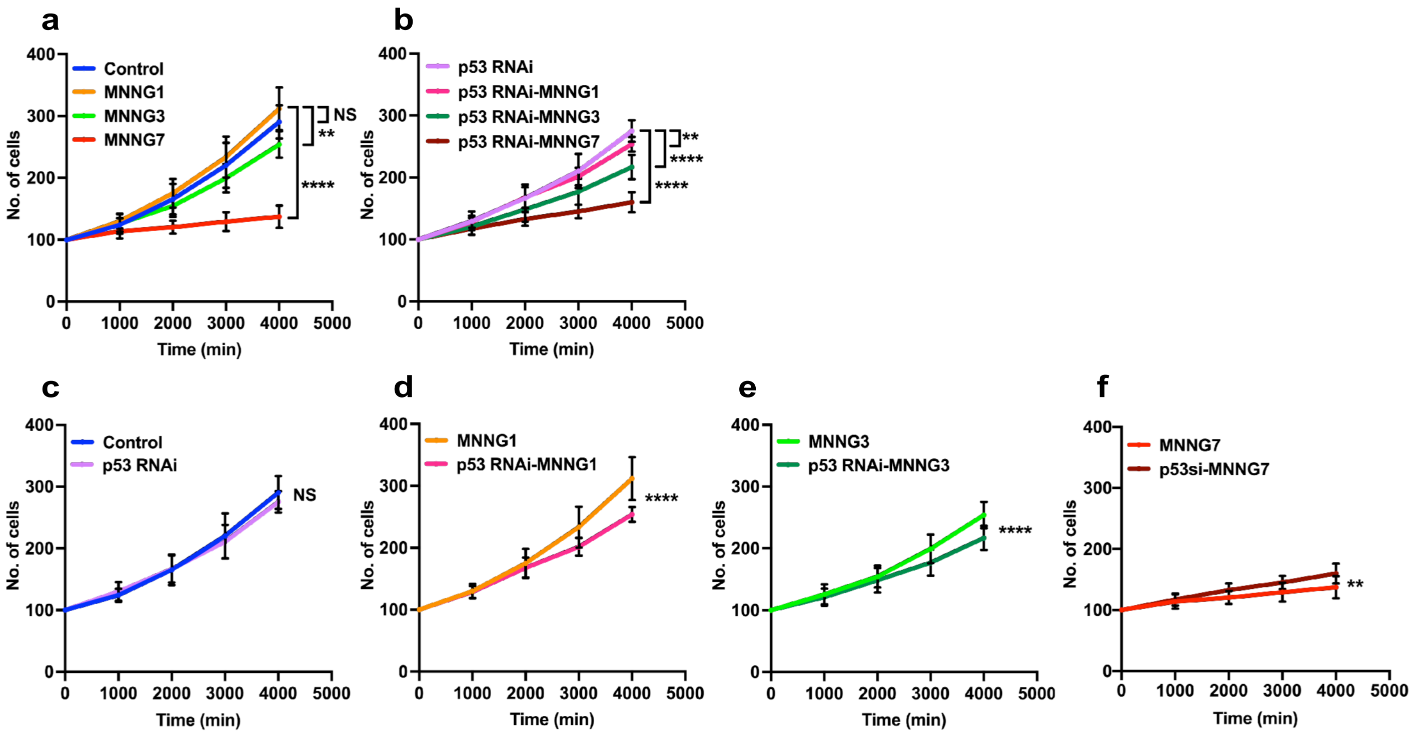


Single-cell tracking

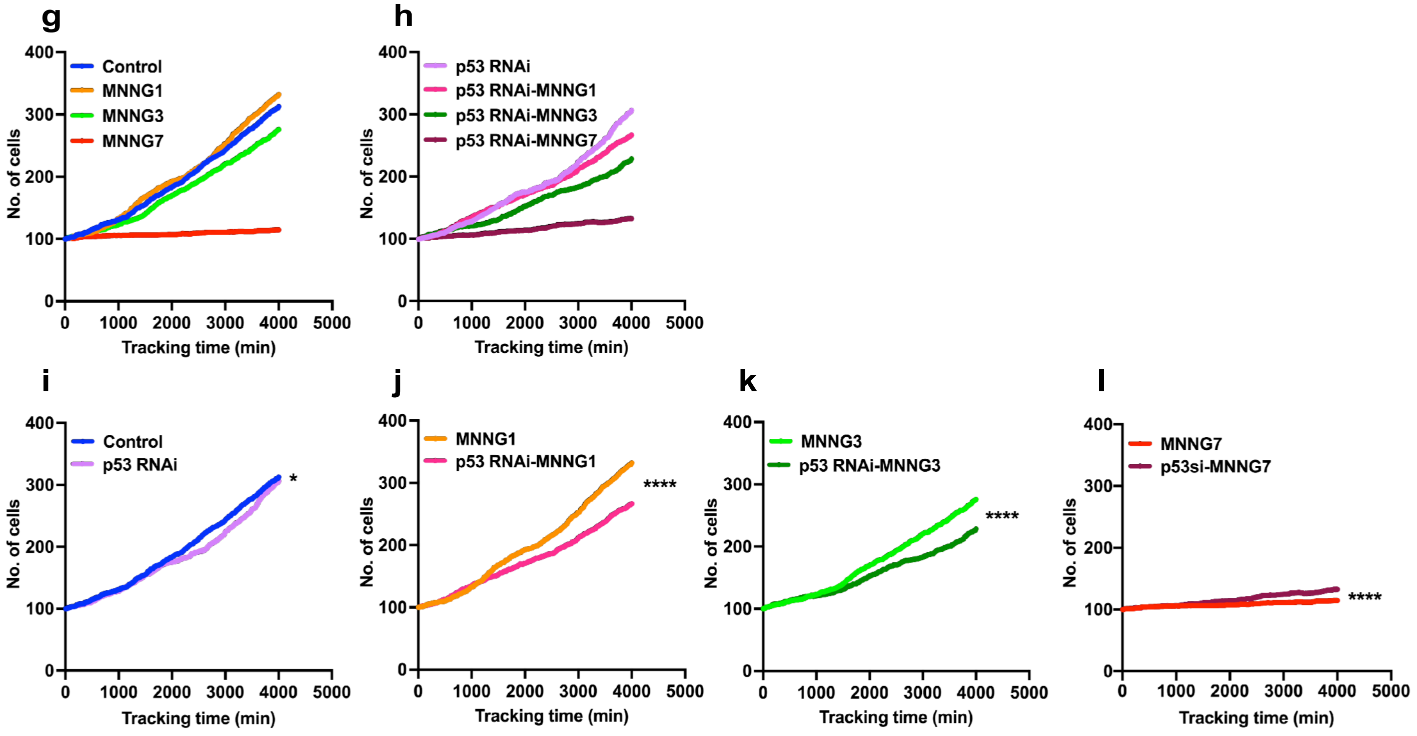


Rancourt et al.

### Figure 2-figure suplement 1

##### Figure 2-figure supplement 1

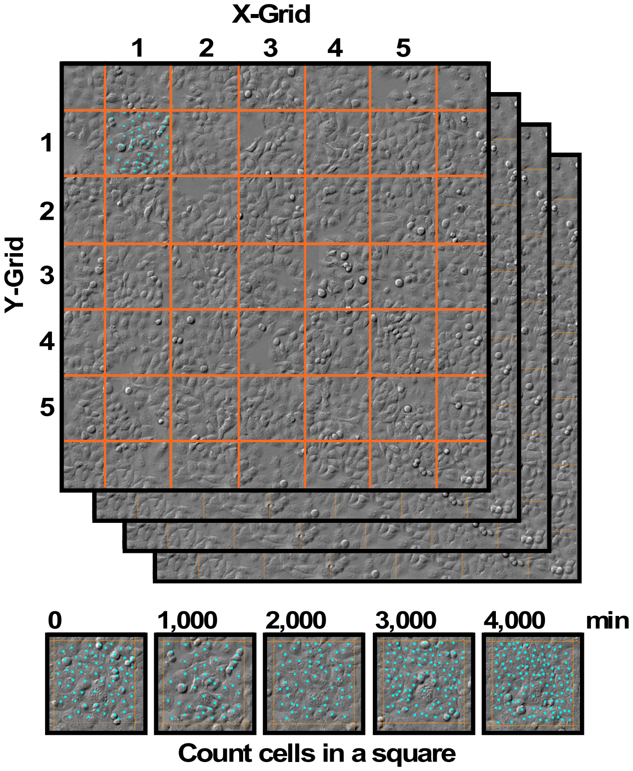


Rancourt et al.

### Figure 3

#### Figure 3

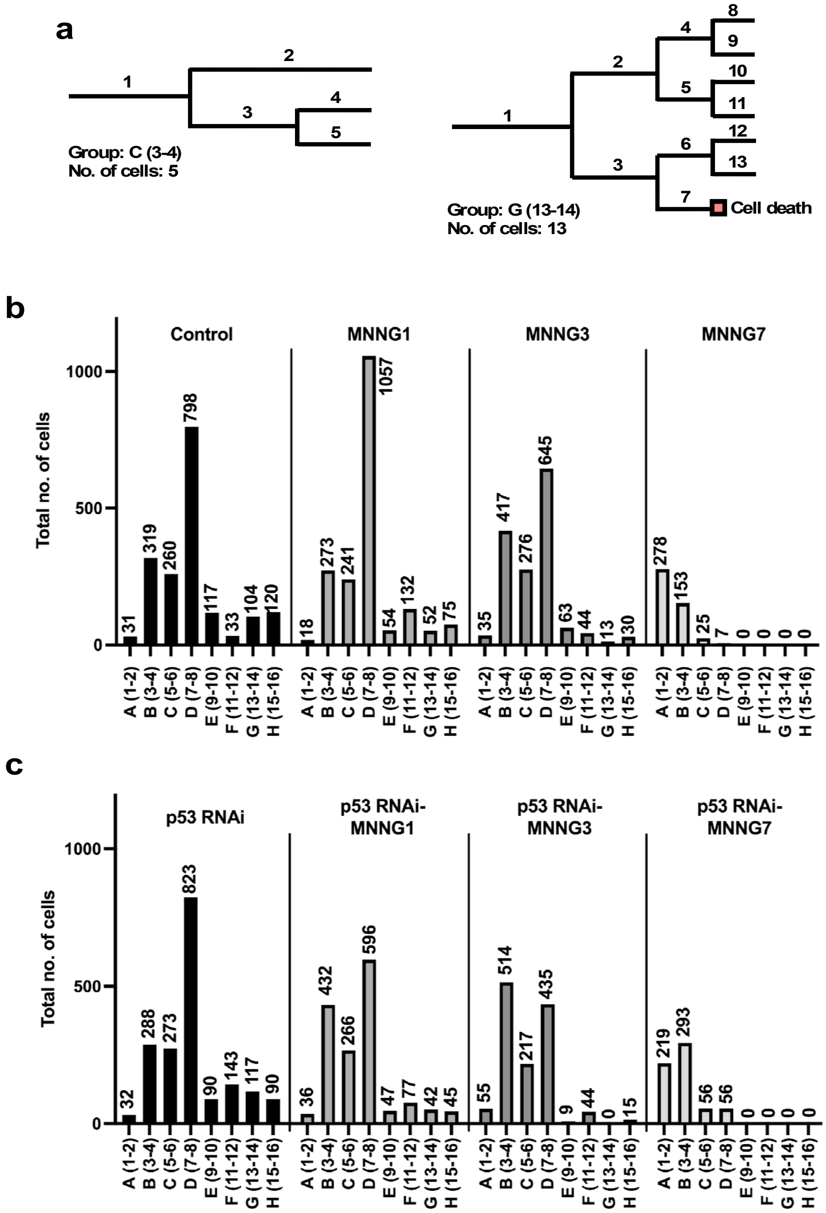


Rancourt et al.

### Figure 4

#### Figure 4

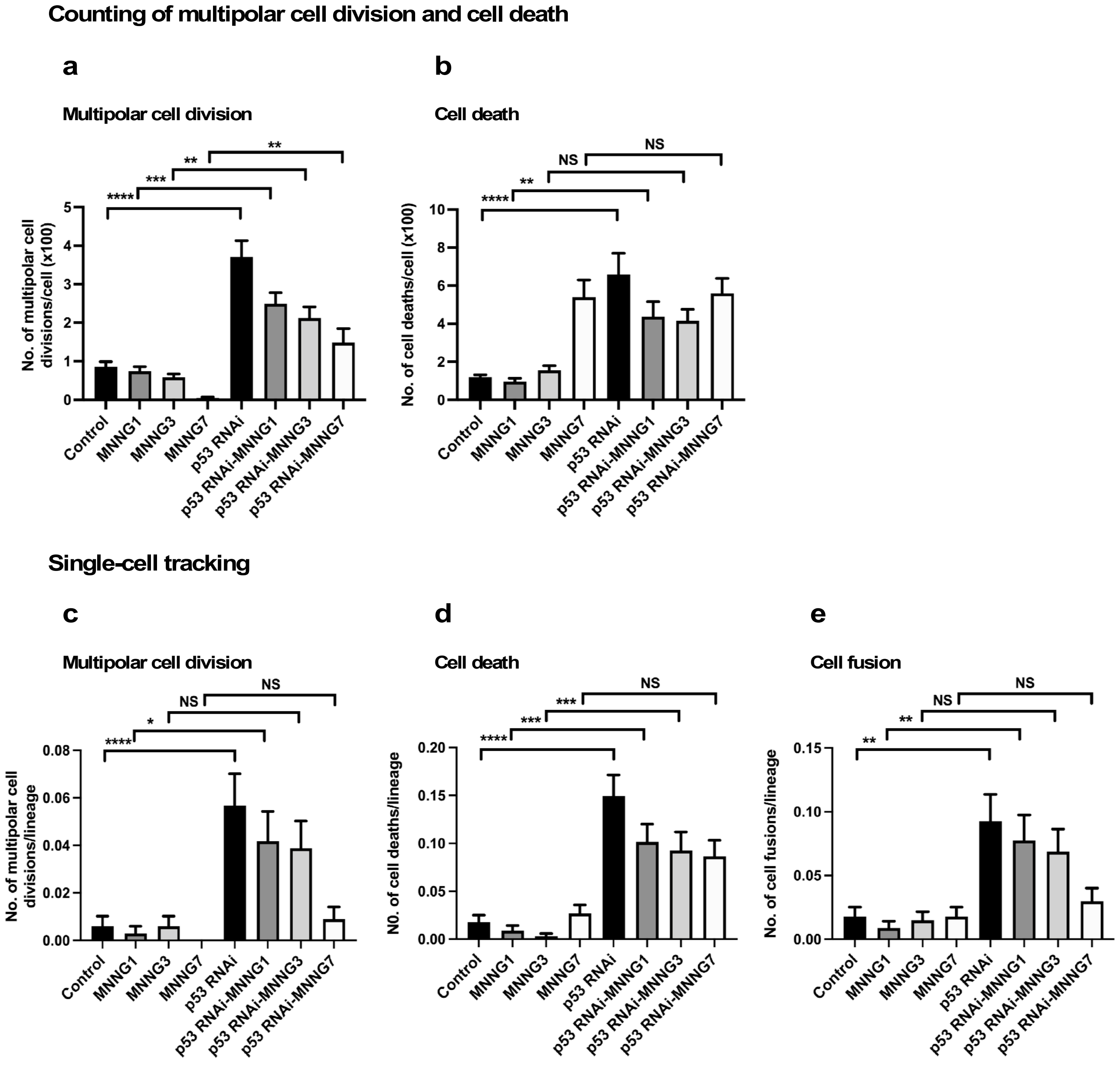


Rancourt et al.

### Figure 4-figure suplement 1

#### Figure 4-figure supplement 1

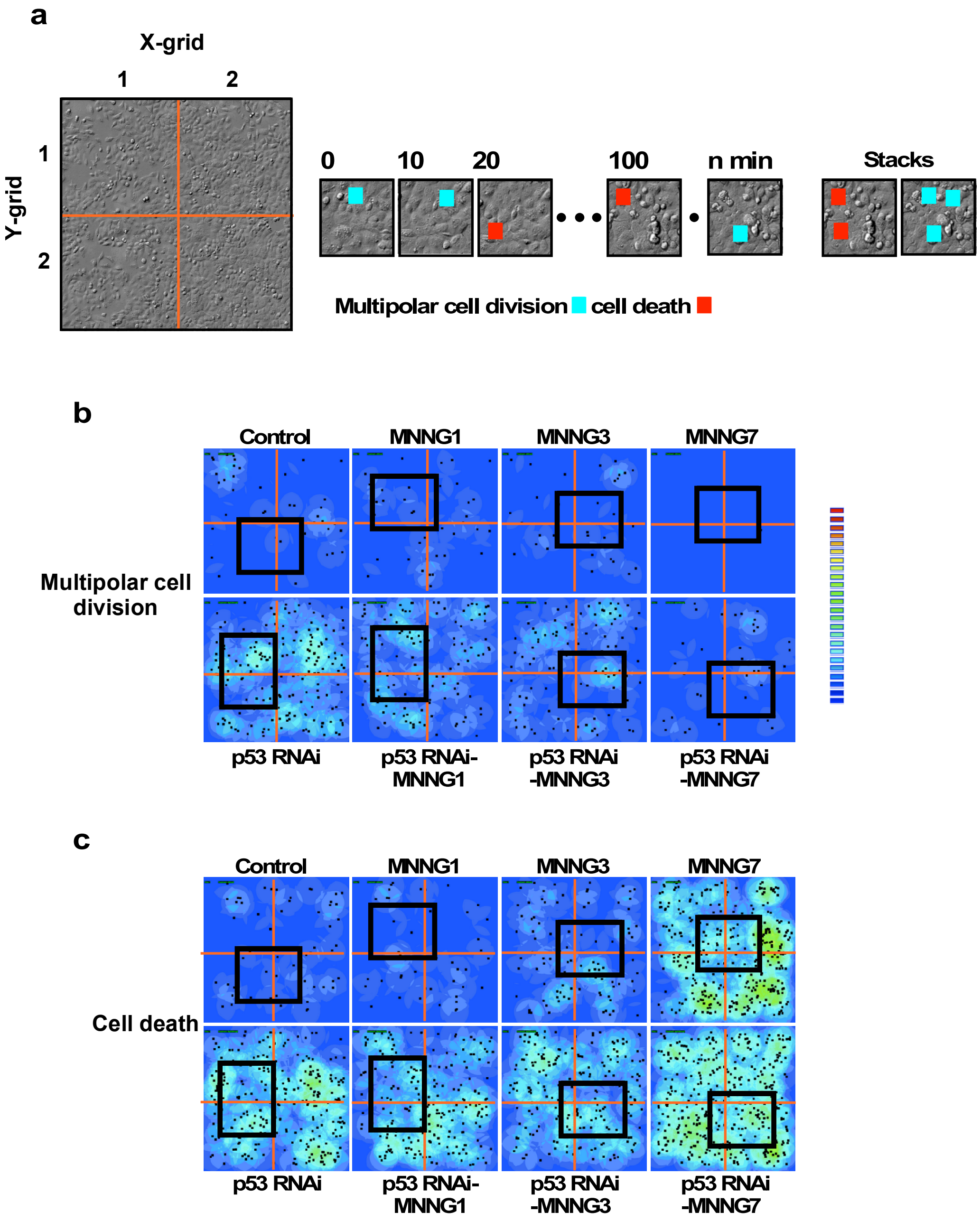

### Figure 5

#### Figure 5

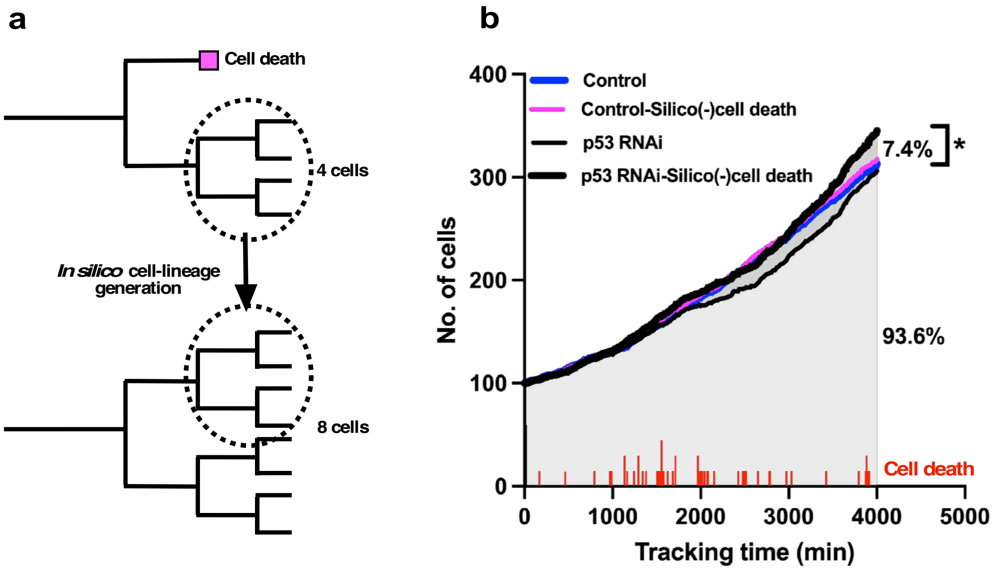


Rancourt et al.

### Figure 7

#### Figure 7

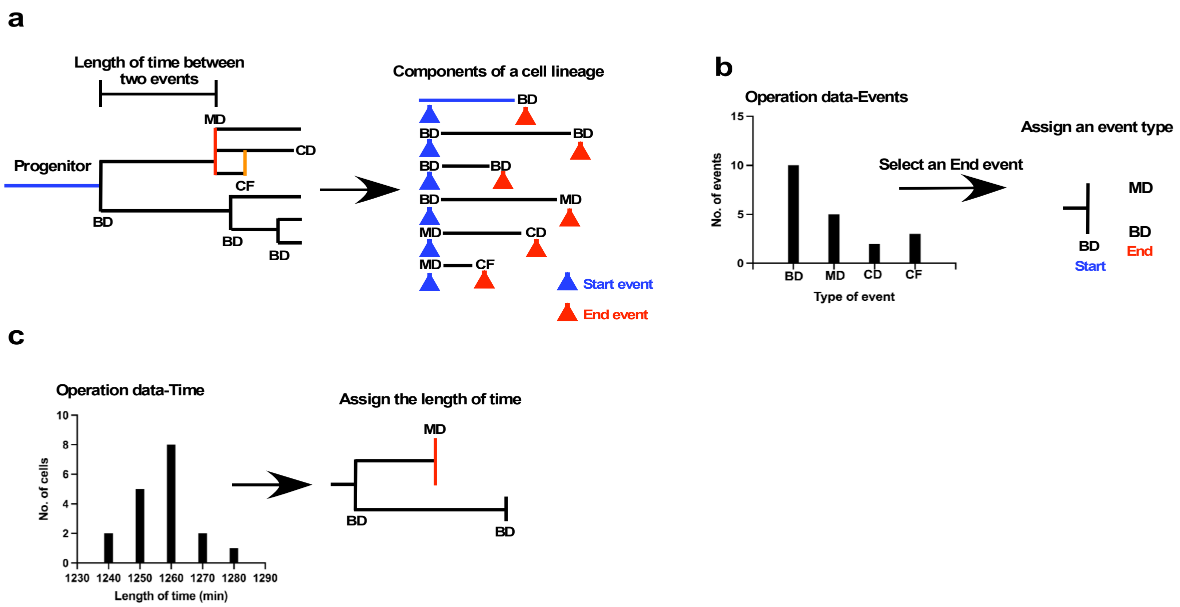


Rancourt et al.

### Figure 7-figure suplement 2-1

##### Figure 7-figure supplement 2

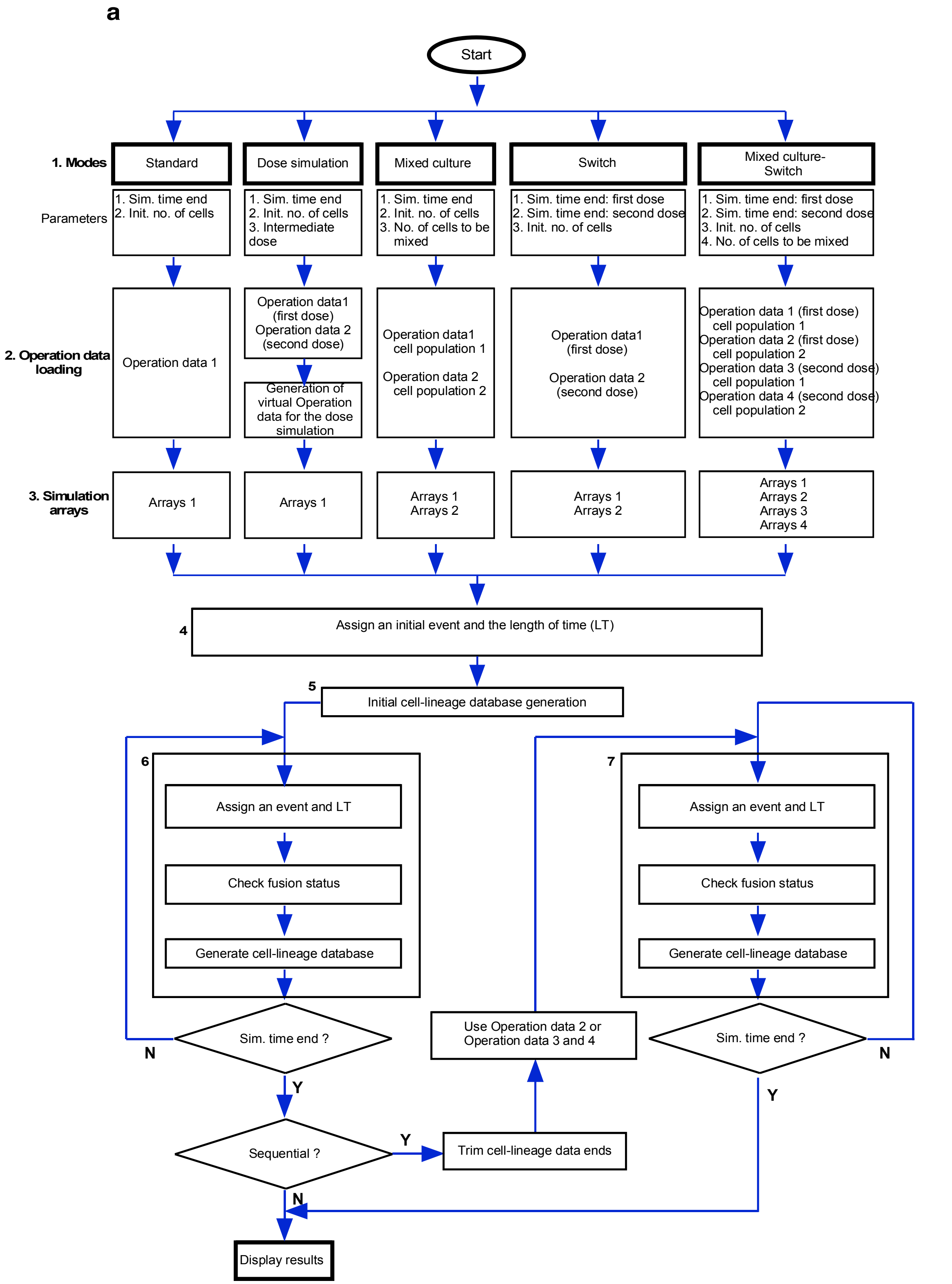


Rancourt et al.

### Figure 7-figure suplement 2-2

#### Figure 7-figure supplement 2

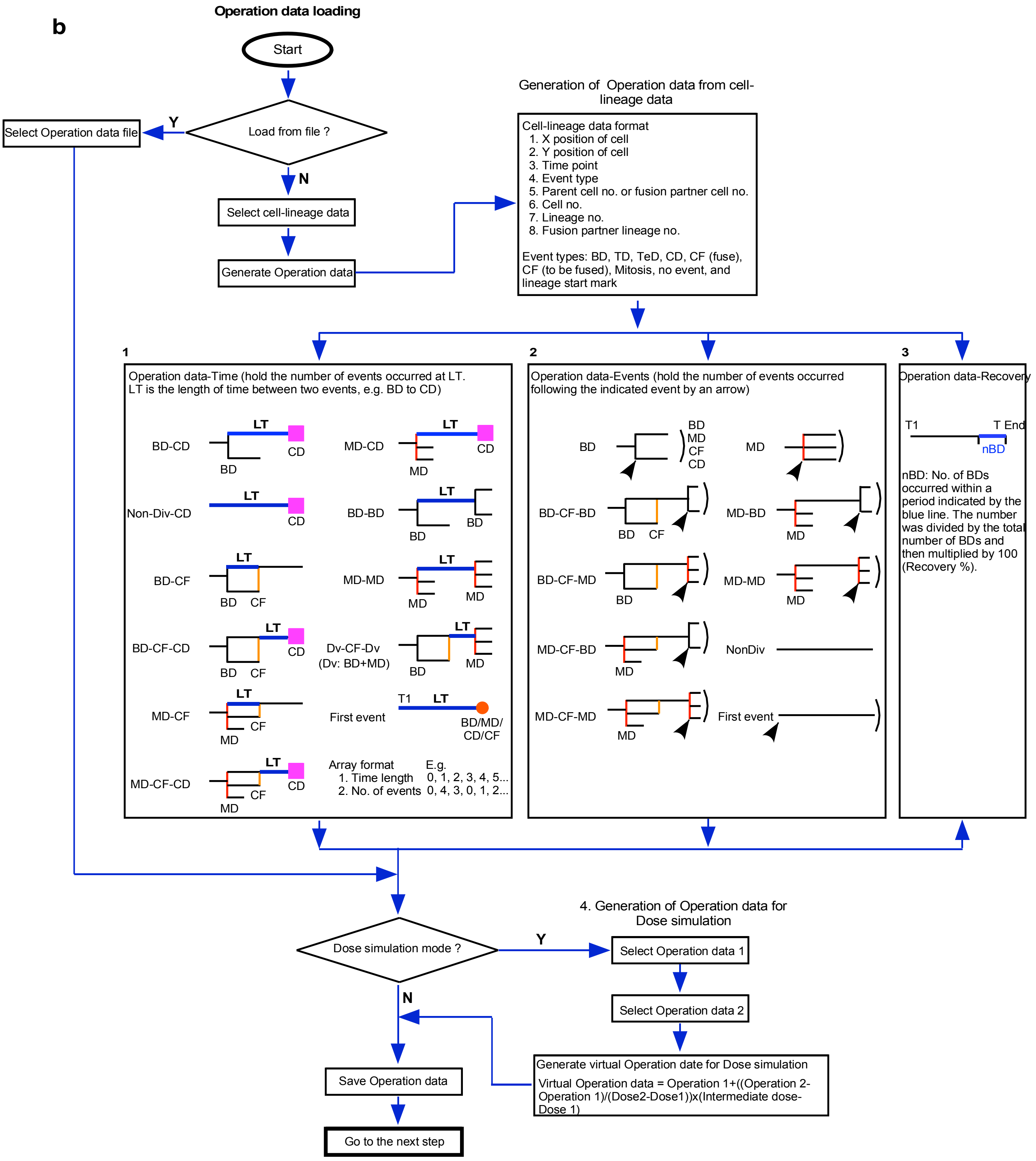


Rancourt et al.

### Figure 7-figure suplement 2-3

#### Figure 7-figure supplement 2

c

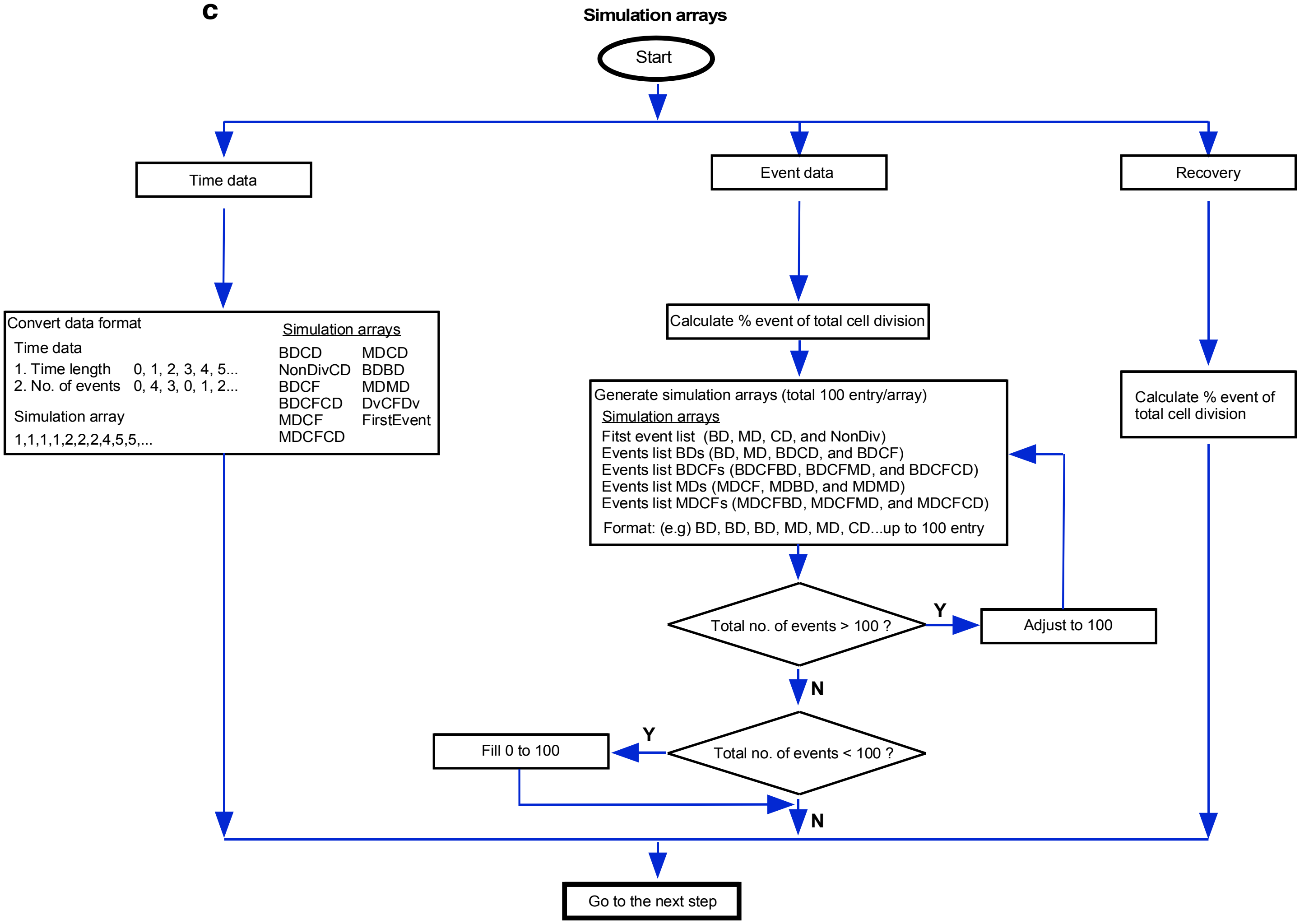


Rancourt et al.

### Figure 7-figure suplement 2-4

#### Figure 7-figure supplement 2

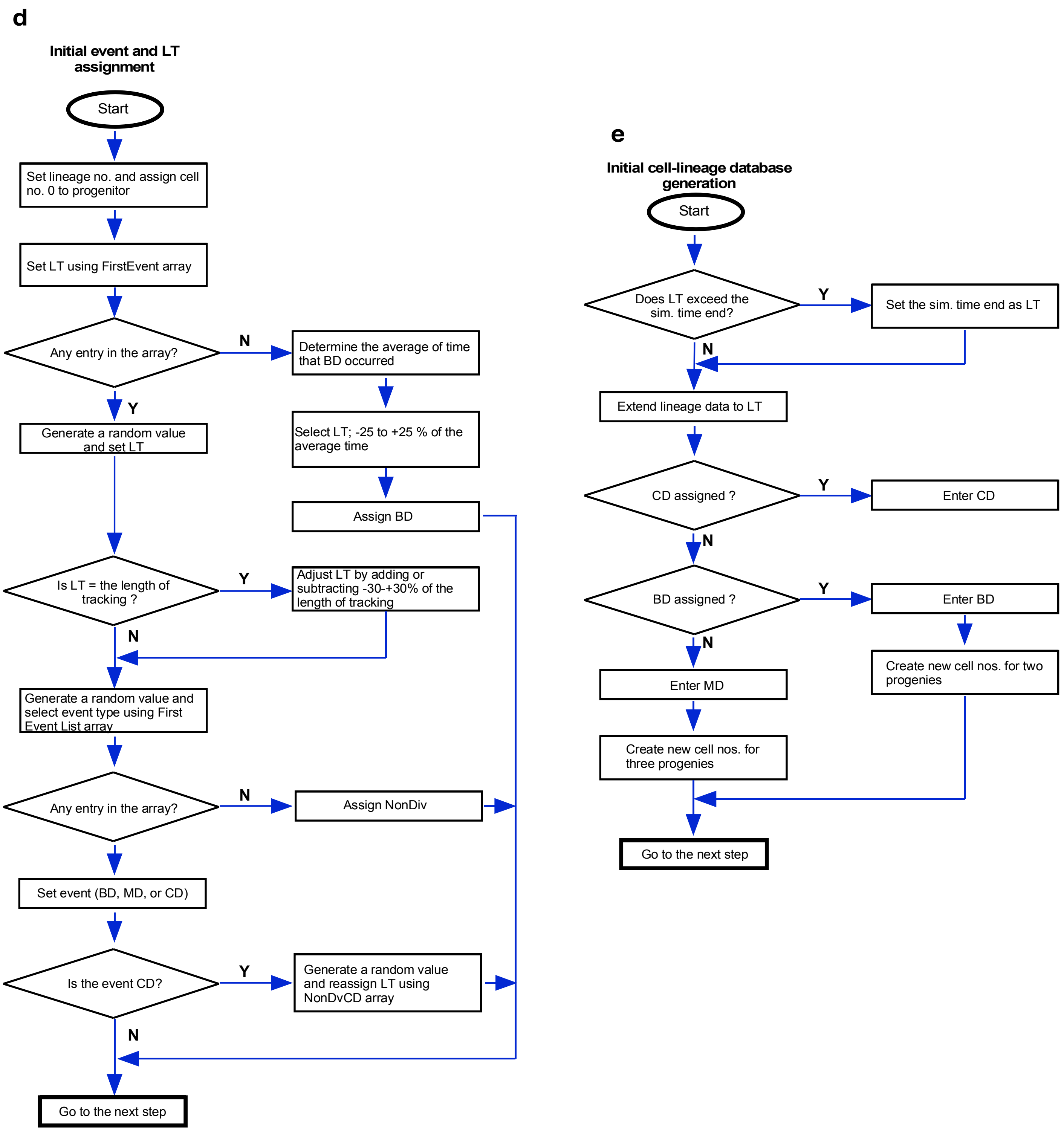


Rancourt et al.

### Figure 7-figure suplement 2-5

#### Figure 7-figure supplement 2

**f**

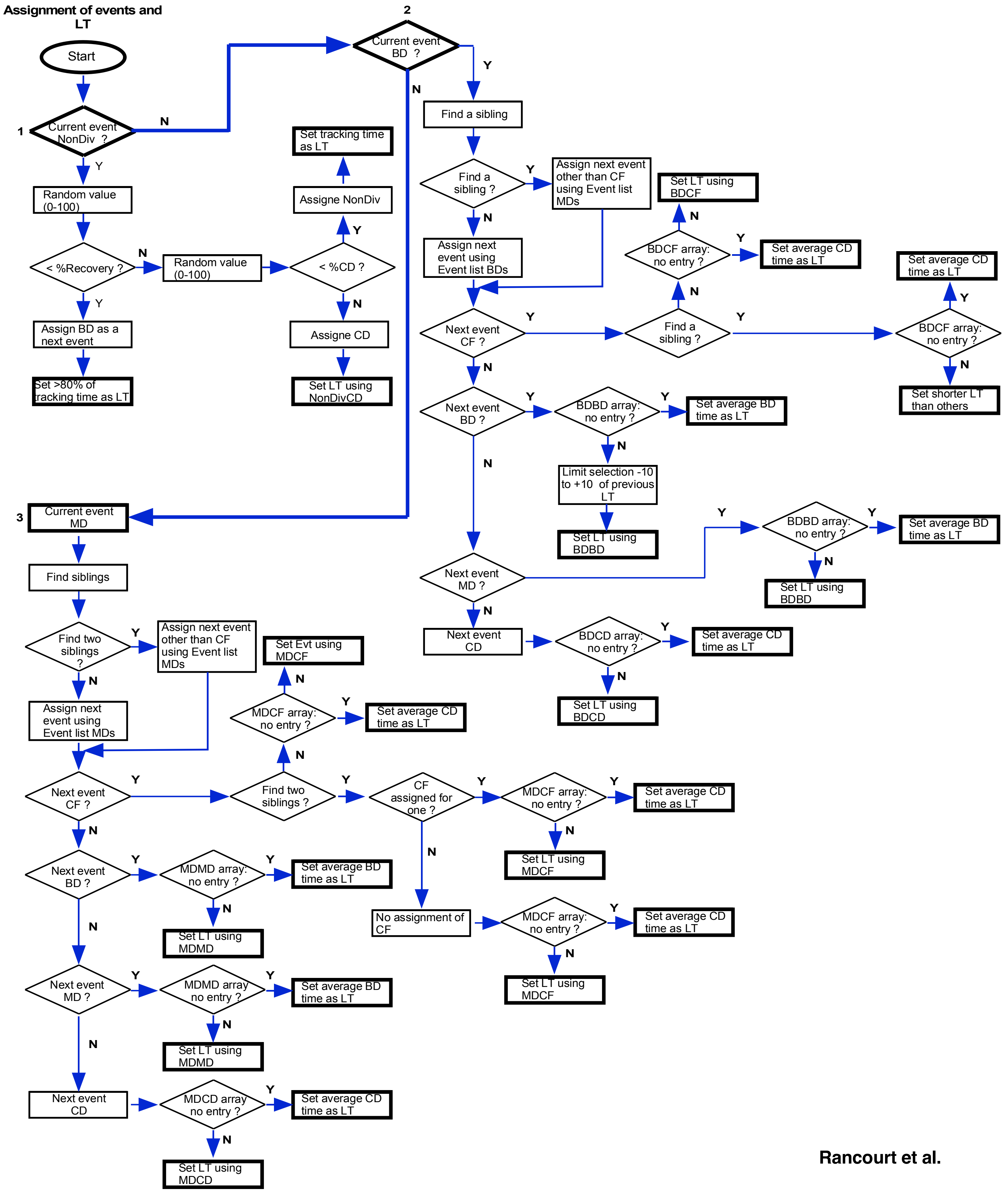


Rancourt et al.

### Figure 8

#### Figure 8

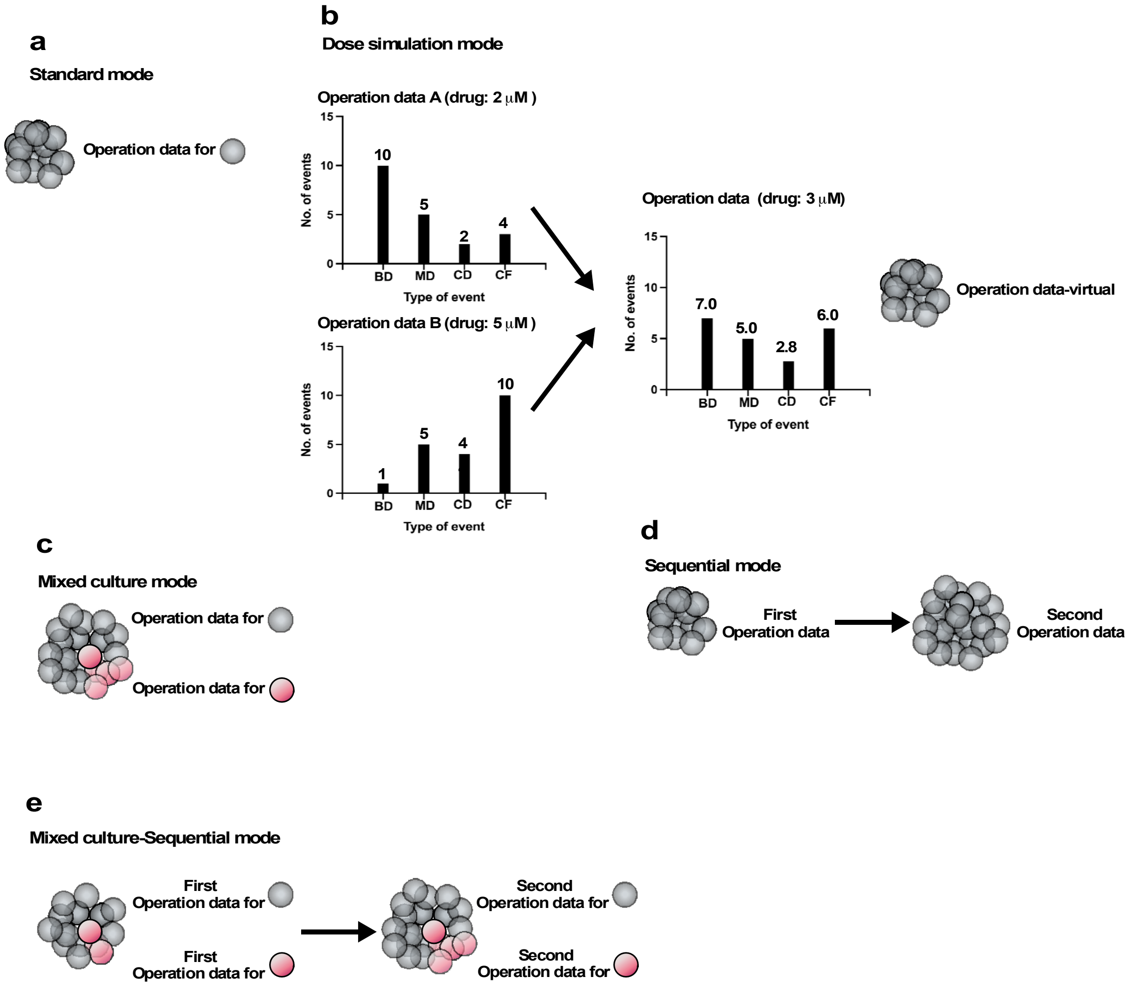


Rancourt et al.

### Figure 9

##### Figure 9

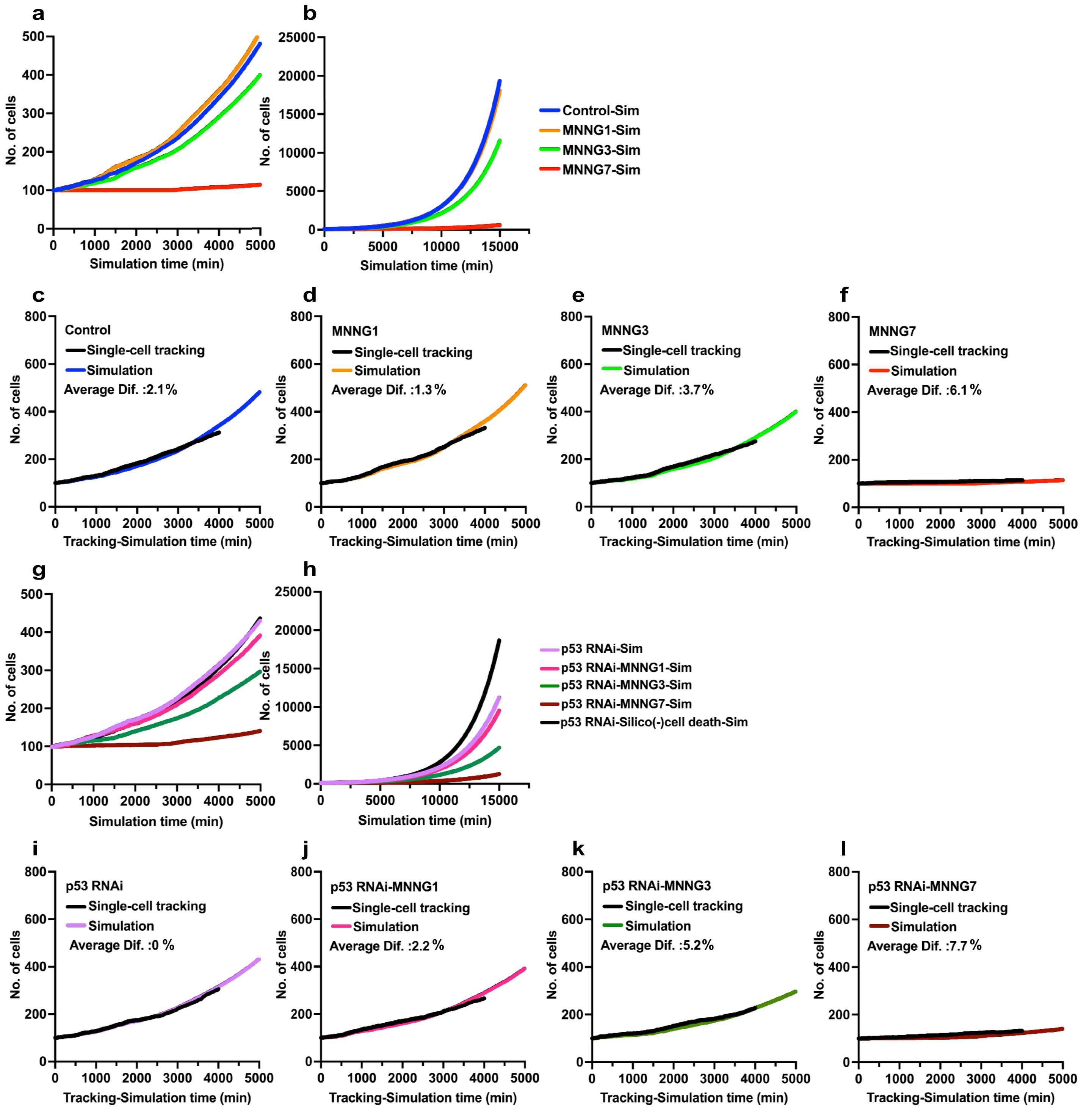


Rancourt et al.

### Figure 10

#### Figure 10

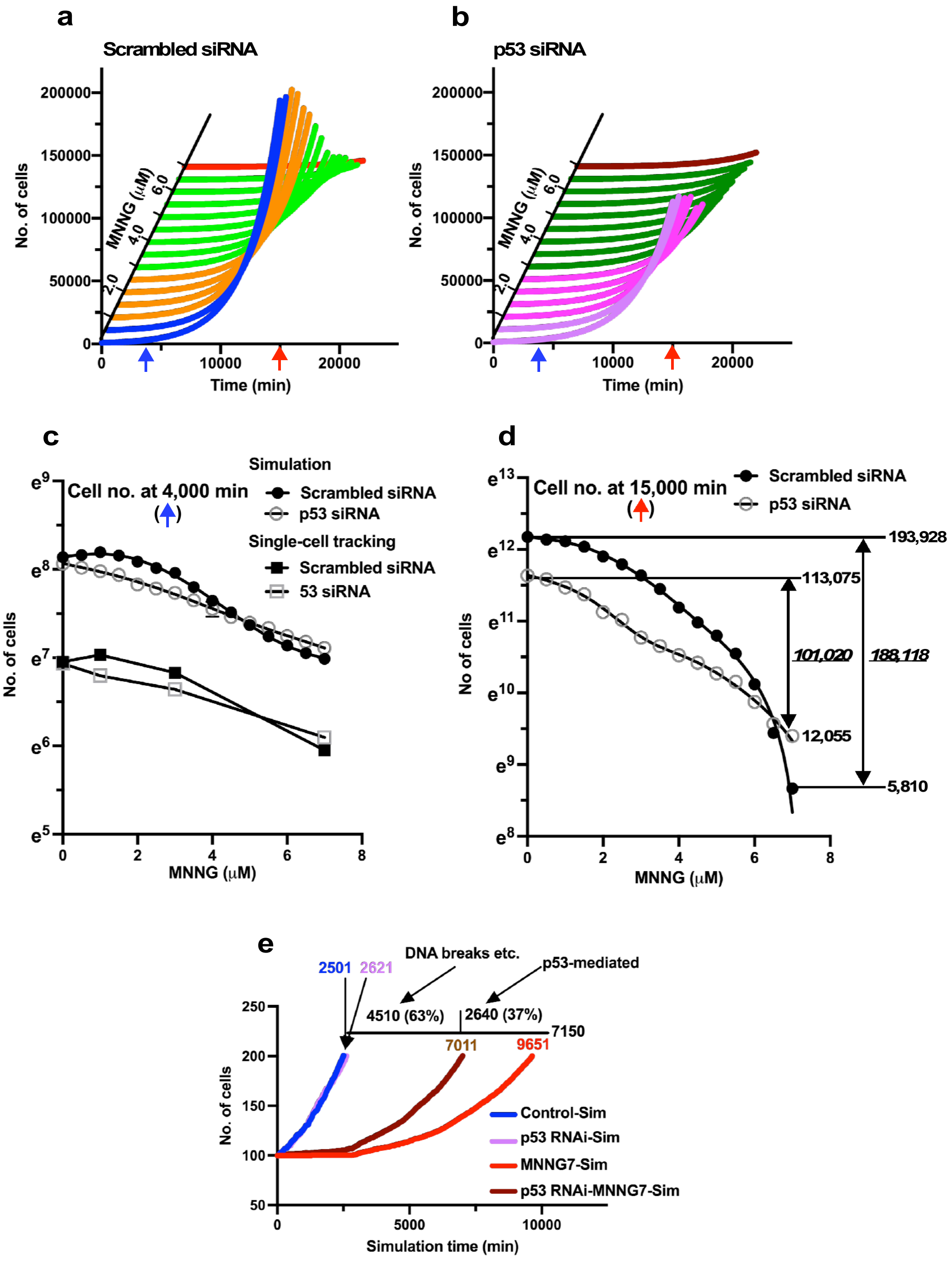


Rancourt et al.

### Figure 11

#### Figure 11

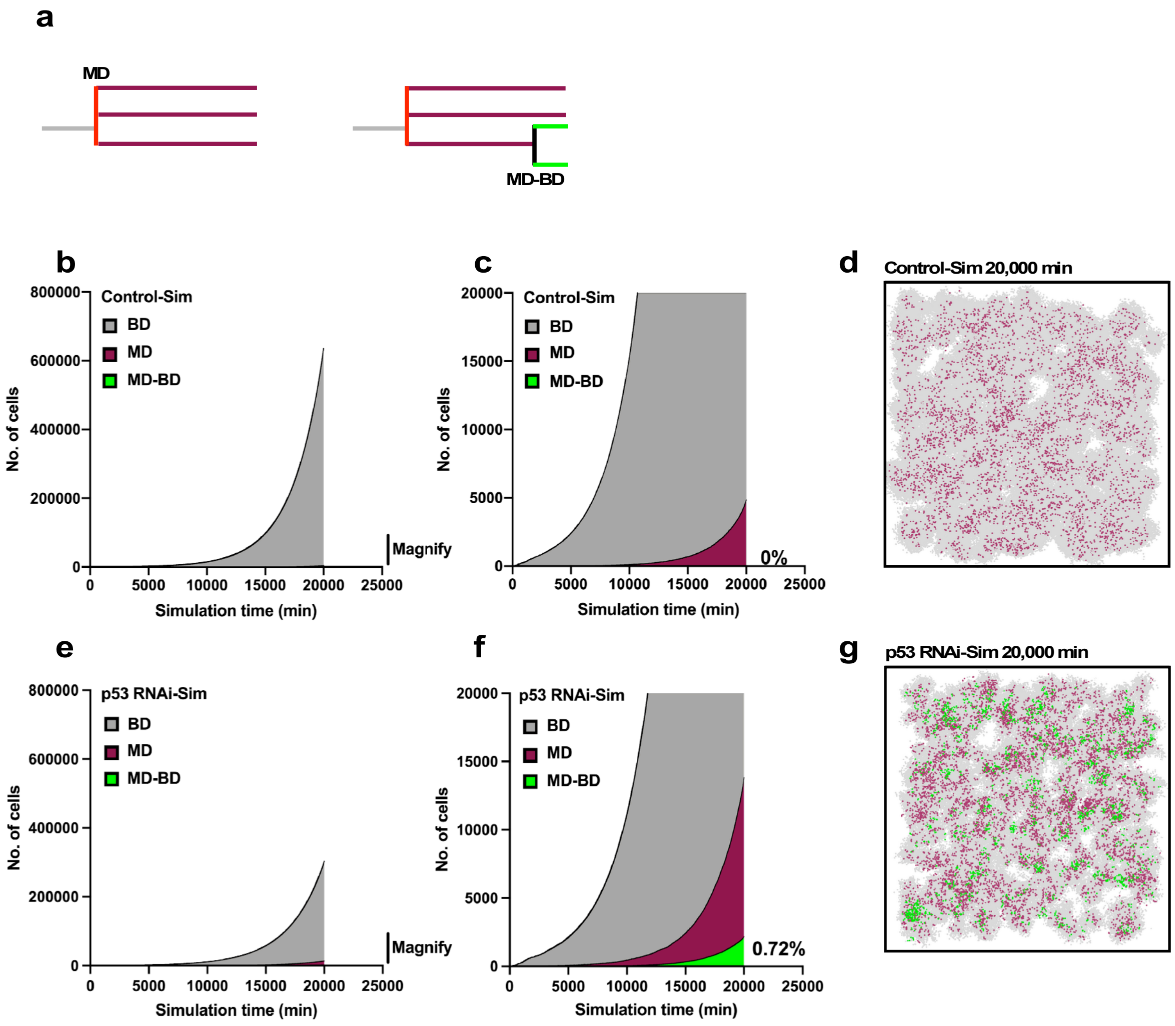


Rancourt et al.

### Figure 12

#### Figure 12

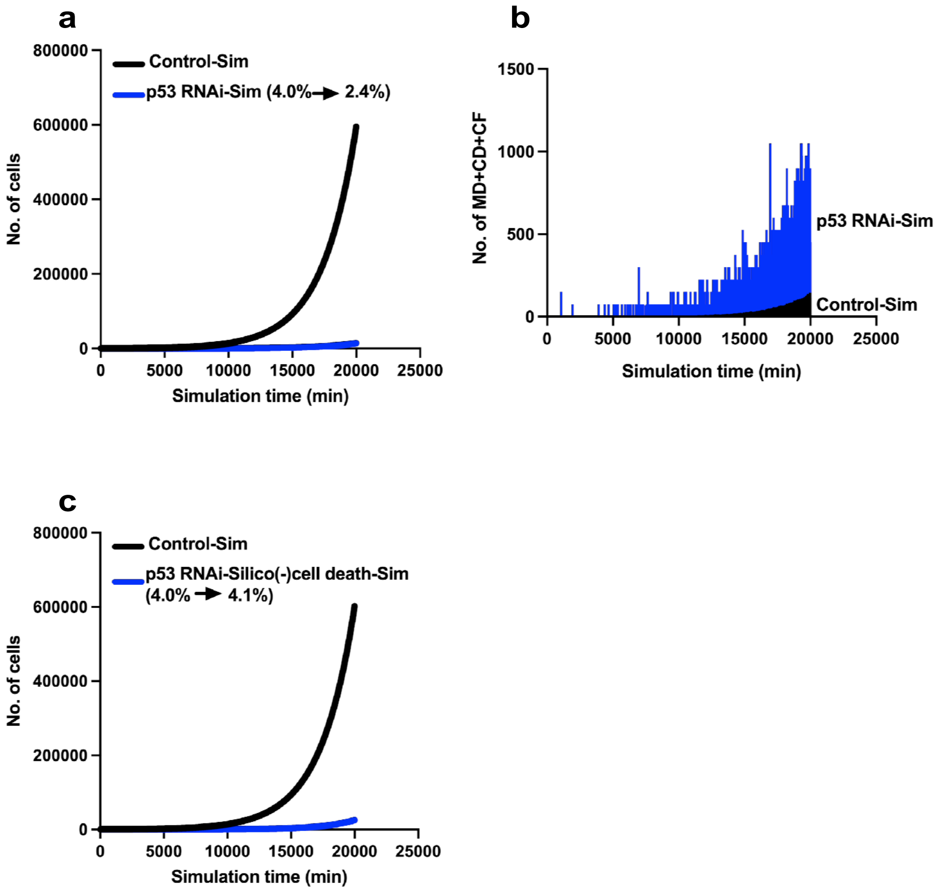


Rancourt et al.

### Figure 13

#### Figure 13

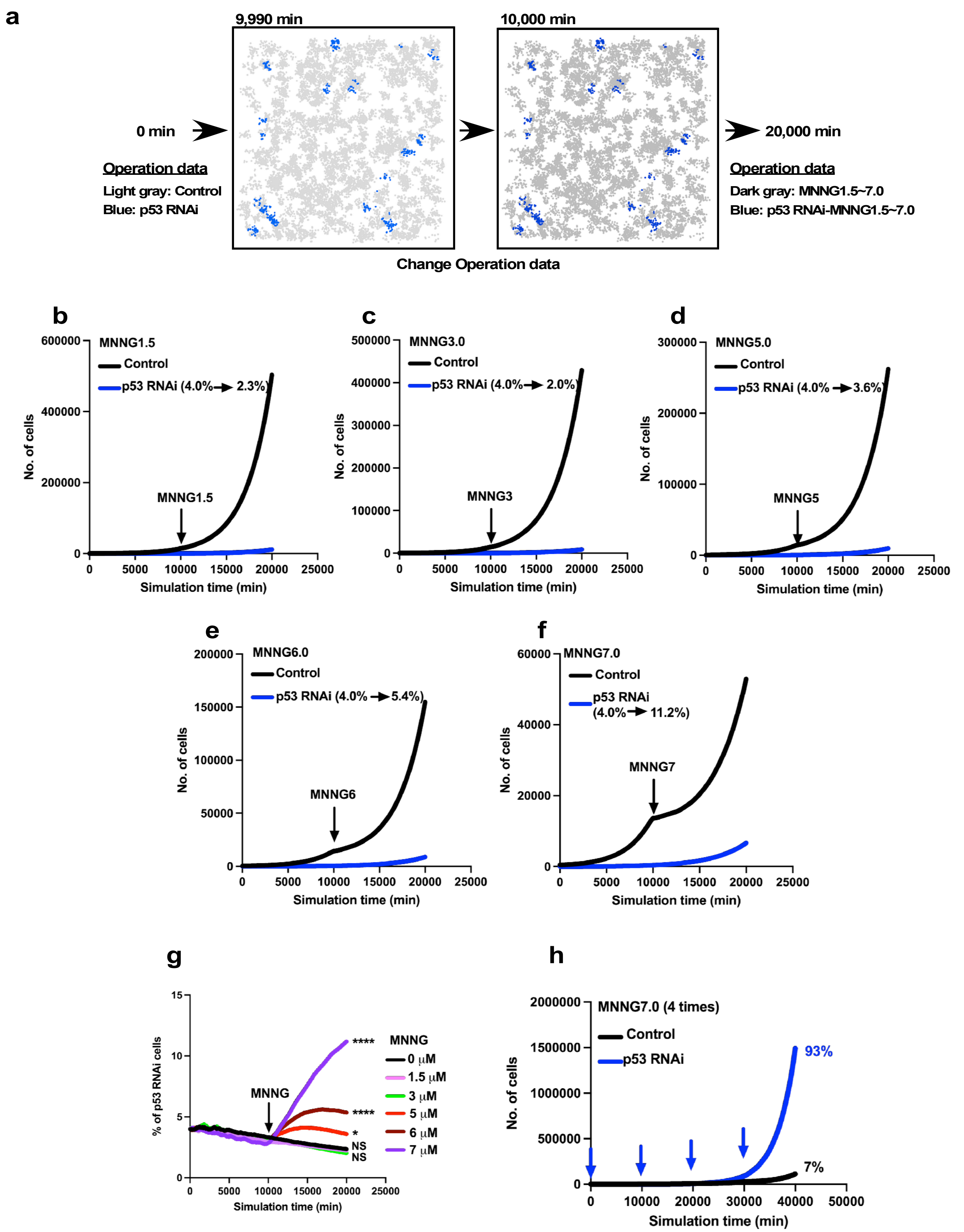


Rancourt et al.

### Figure 14

#### Figure 14

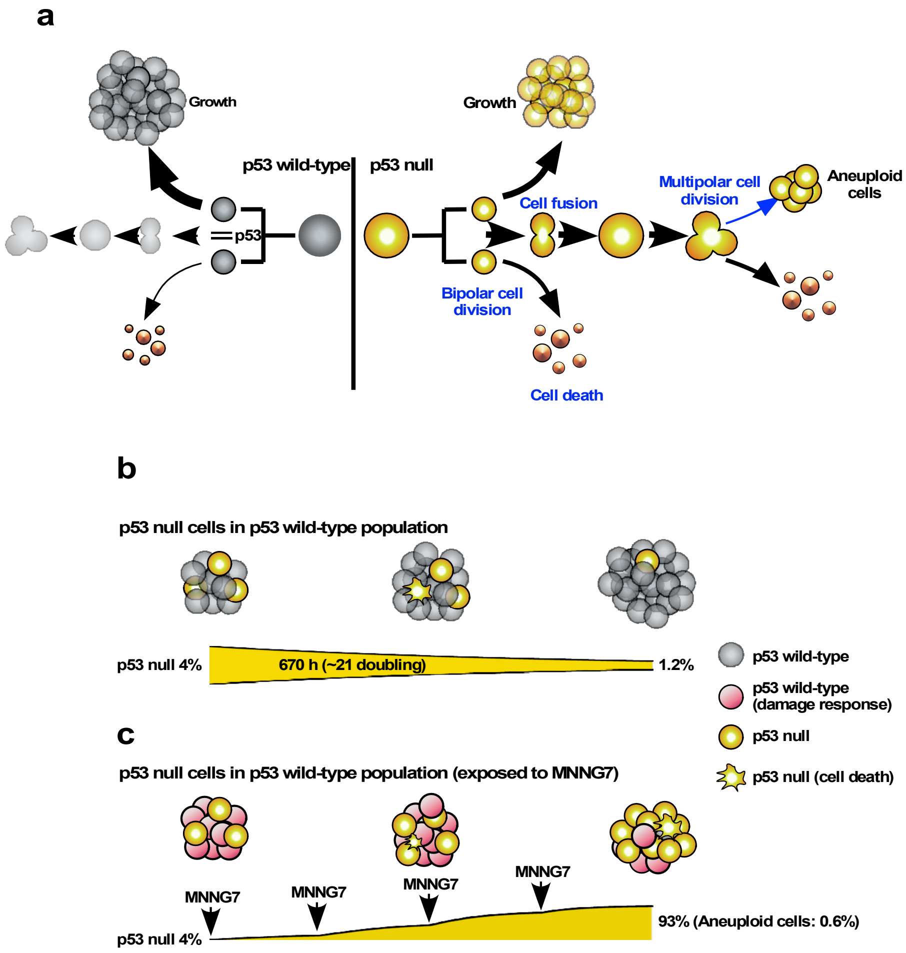


Rancourt et al.
