## Supplementary material for "Empirical single-cell tracking and cell-fate simulation reveal dual roles of p53 in tumor suppression": Figure 6

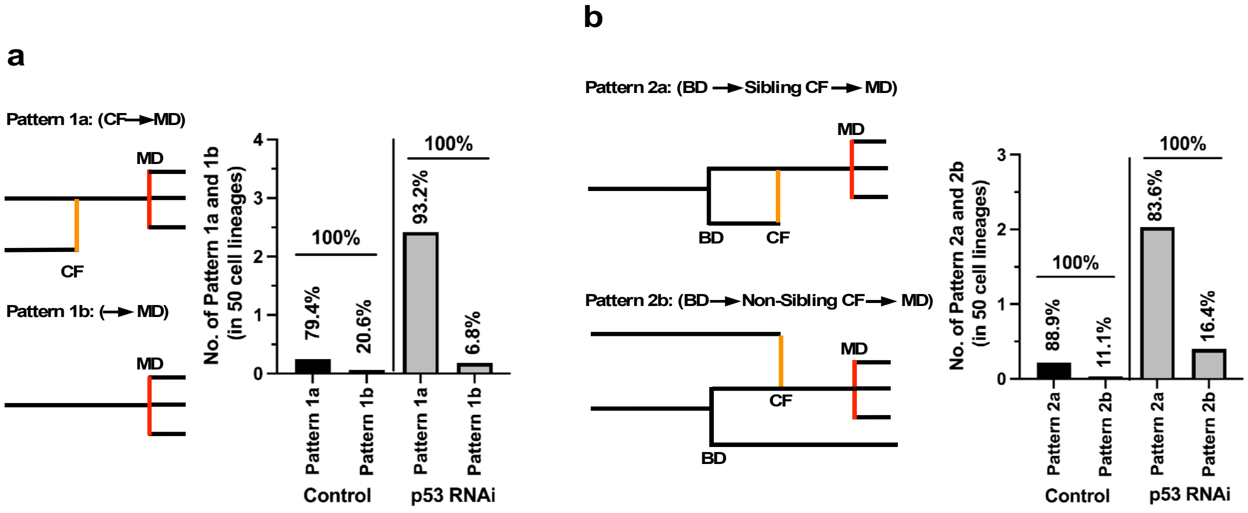


c

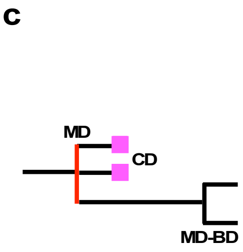


| | Reproductive progeny (MD-BD) | |
|---|---|---|
| | MD-BD/100 lineages | % of the progeny of MD |
| Control | 0.06 | 3.22 |
| p53 RNAi | 0.61 | 3.63 |

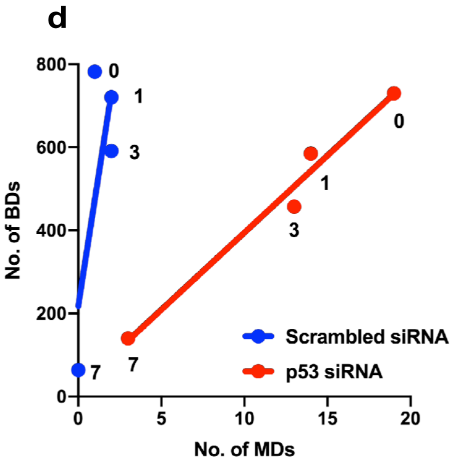


Rancourt et al.
