## Supplementary material for "Empirical single-cell tracking and cell-fate simulation reveal dual roles of p53 in tumor suppression": Figure 7-figure suplement 1

### Figure 7-figure supplement 1

**a**

Begin:

**Create live cell imaging videos**

**Segment images**

**Perform automatic single-cell tracking and create cell-lineage database**
    Verify tracking data manually
    Restart automatic single-cell tracking

        Record the position of each cell, an event type that occurred in a cell, and the relation of a cell to other cells (parent and siblings)

**Display cell-lineage maps**

**Analyze tracking data**

**Create Operation data for simulation**

**Save Operation data**

End:

**b**

Begin:

**Upload Operation data from a saved file**
    If no file was found, create Operation data from a cell-lineage database

**Set simulation parameters**
    Set a simulation time
    Set the number of progenitors

**Create Operation arrays**
    Create an array: the First event Operation data-Events
    Create an array: the First event-Operation data-Time
    Create arrays: Operation data-Events for other classified event patterns
    Create arrays: Operation data-Time for other classified event patterns

**Create an array that holds cell-lineage data**

**Assign an event and the length of time to progenitors**
    Assign a cell-lineage number to each progenitor
    Assign cell number 0 to progenitors
    Assign an event type to each progenitor
        Select an event type by using the First event-Operation data-Events
    Assign the length of time to each progenitor
        Select the length by using the First event-Operation data-Time

**Assign an event type and the length of time to progeny**
    For x = simulation time
        *Create a temporary array that holds cell-lineage data*

        *Assign an event type to a progeny using relevant Operation data-Events*
            Select an event type

        *Assign the length of time to a progeny using relevant Operation data-Time*
            Select the length of time

                If bipolar cell division is assigned, adjust the length of time within +- 10% of its parent cell

                If bipolar or multipolar cell divisions are assigned, generate the relevant number of progeny (2 and 3 for bipolar and multipolar cell division, respectively)
                    Assign a new cell number

        *Save assigned data into the temporary array*

        *Transfer the temporarily saved data to the cell-lineage data array*
            If the size of the cell-lineage data array exceeds the allocated size, expand the array size

        *Clear the temporary array*

    Repeat

**Display cell-lineage maps**
**Generate animation**

**Clear all arrays**

End:

Rancourt et al.
