## Supplementary material for "Empirical single-cell tracking and cell-fate simulation reveal dual roles of p53 in tumor suppression": Figure 7-figure suplement 2-6

### Figure 7-figure supplement 2

g

**Checking fusion status**

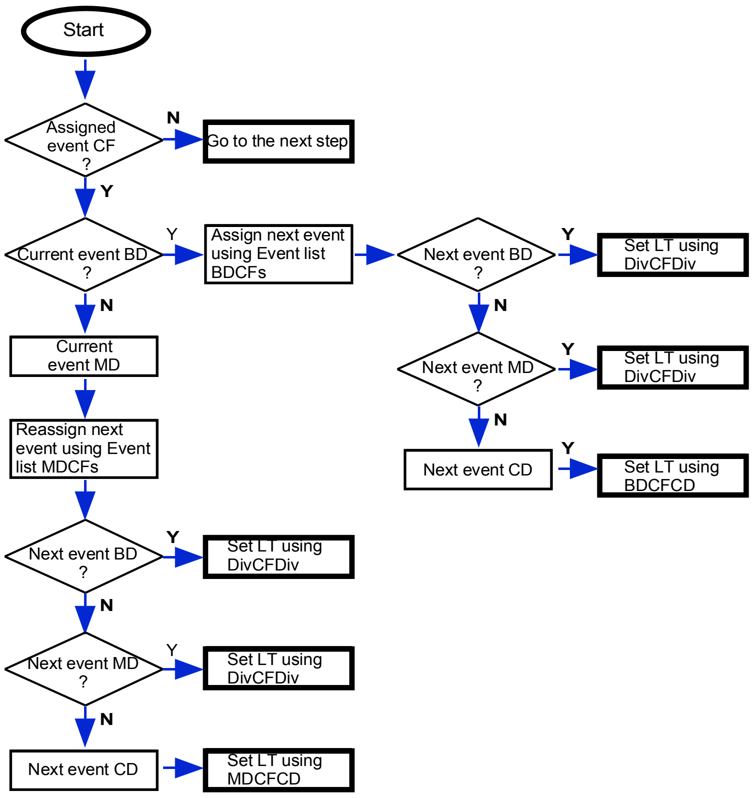


h

**Lineage database generation**

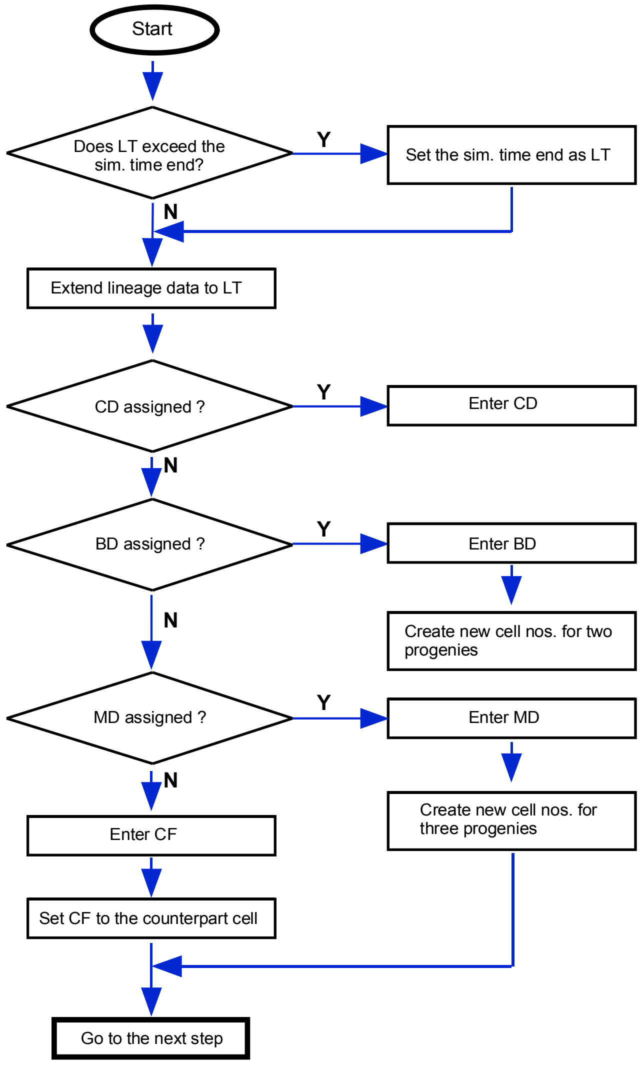


i

**Trimming of cell lineage data ends**

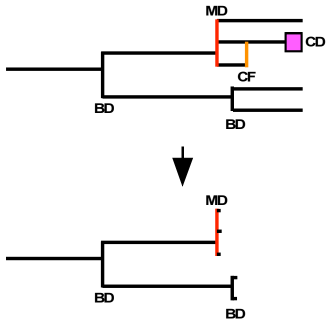


Rancourt et al.
